# Purkinje cell outputs selectively inhibit a subset of unipolar brush cells in the input layer of the cerebellar cortex

**DOI:** 10.1101/2021.03.25.437031

**Authors:** Chong Guo, Stephanie Rudolph, Morgan E. Neuwirth, Wade G. Regehr

**Affiliations:** Department of Neurobiology, Harvard Medical School, 220 Longwood Ave, Boston MA 02115

## Abstract

Circuitry of the cerebellar cortex is regionally and functionally specialized. Unipolar brush cells (UBCs), and Purkinje cell (PC) synapses made by axon collaterals in the granular layer, are both enriched in areas that control balance and eye-movement. Here we find a link between these specializations: PCs preferentially inhibit mGluR1-expressing UBCs that respond to mossy fiber inputs with long lasting increases in firing, but PCs do not inhibit mGluR1-lacking UBCs. PCs inhibit about 29% of mGluR1-expressing UBCs by activating GABA_A_ receptors (GABA_A_Rs) and inhibit almost all mGluR1-expressing UBCs by activating GABA_B_Rs. PC to UBC synapses allow PC output to regulate the input layer of the cerebellar cortex in diverse ways. GABA_A_R-mediated feedback is fast, unreliable, noisy, and suited to linearizing input-output curves and decreasing gain. Slow GABA_B_R-mediated inhibition allows elevated PC activity to sharpen the input-output transformation of UBCs, and allows dynamic inhibitory feedback of mGluR1-expressing UBCs.

## Introduction

Although different lobules of the cerebellar cortex are engaged in diverse motor and non-motor behaviors (Ito, 1998; J. J. Kim & Thompson, 1997; Massion, 1992; Murdoch, 2010; Schmahmann & Sherman, 1998; Tereszko & Dudek, 2014; Van Overwalle et al., 2014; Villanueva, 2012), a basic circuit motif is repeated within each functional compartment. Mossy fibers (MFs) provide excitatory inputs to granule cells (Delvendahl & Hallermann, 2016; DiGregorio et al., 2002; Huang et al., 2013; Sotelo, 2008), which in turn excite Purkinje cells (PCs) that provide the output of the cerebellar cortex. In addition, Golgi cells (GoCs) inhibit granule cells (GrCs), and molecular layer interneurons (MLIs) inhibit PCs (Eccles, 2013). Regional synaptic and cellular specializations exist beyond this basic circuit motif, presumably to meet the computational demands associated with specific behaviors (DiÑO et al., 1999; Guo et al., 2016; Kozareva et al., 2020; Sekerková et al., 2014; Suvrathan et al., 2016).

One of the most obvious specializations is that the density of unipolar brush cells (UBCs) exhibits large regional variation (DiÑO et al., 1999; Sekerková et al., 2014). UBCs are excitatory interneurons located in the granular layer that are usually innervated by a single MF (Enrico Mugnaini & Floris, 1994). UBCs contribute to temporal processing by converting short-lived MF signals into long-lasting changes in (Kennedy et al., 2014; Kinney et al., 1997; Kreko-Pierce et al., 2020; Locatelli et al., 2013; Enrico Mugnaini & Floris, 1994; E. Mugnaini et al., 2011; Rossi et al., 1995; van Dorp & De Zeeuw, 2014). UBCs. MFs excite UBCs that express metabotropic glutamate receptor type I (mGluR1), suppress firing in UBCs where mGluR2 is prominent, and evoke more complex responses in other UBCs that express both mGluR1 and mGluR2 (Borges-Merjane & Trussell, 2015; Guo et al., 2020; Kinney et al., 1997; Knoflach & Kemp, 1998; Rossi et al., 1995; Russo et al., 2008; van Dorp & De Zeeuw, 2014; Zampini et al., 2016). UBCs are often disregarded in descriptions of the circuitry of the cerebellar cortex. However, UBCs are present in all areas and are exceptionally dense in some regions, notably those involved in vestibular function (DiÑO et al., 1999; Takács et al., 1999). UBCs are also present in larger mammals including humans (DiÑO et al., 1999; Munoz, 1990), where their high densities and widespread distributions suggest they may have a special role in cerebellar computations relating to higher cognition.

PC feedback is another noteworthy regional specialization. In addition to sending an axon to the deep cerebellar nuclei, each PC axon has a collateral that inhibits PCs (Bernard & Axelrad, 1993; Bernard et al., 1993; Bornschein et al., 2013; Orduz & Llano, 2007; Watt et al., 2009; Witter et al., 2016) and several types of inhibitory interneurons in the cerebellar cortex (Crook et al., 2007; Hirono et al., 2012; Witter et al., 2016). In some regions, PCs make extensive contacts within the granular layer and inhibit GrCs (Guo et al., 2016). This feedback is mediated primarily by GABA_A_ receptors, and has a prominent slow component that suggests the involvement of the extrasynaptic high-affinity GABA_A_ receptors present on GrC dendrites. These PC to GrC synapses allow the output of the cerebellar cortex to provide slow negative feedback to inhibit the input layer. Like UBCs these synapses are also most prominent in regions involved in vestibular function.

The regional overlap of high densities of UBCs and granular layer PC synapses raises the possibility that PCs might inhibit UBCs and thereby allow PC feedback to refine temporal processing. Here we examine whether PCs inhibit UBCs. We find that indeed PCs phasically inhibit a subset of UBCs by activating GABA_A_ receptors, and that UBCs with prominent mGluR1 signaling are preferentially targeted. PCs also provide long-lasting inhibition to most mGluR1-expressing UBCs by activating GABA_B_ receptors. This component of inhibition interacts with voltage-gated conductances in UBCs to sharpen the input-output transformation of UBCs. Additionally, slow PC feedback allows PC population activity to directly control spontaneous firing in UBCs. In these ways, PC inhibition of mGluR1-expressing UBCs provides direct feedback from the output layer to input layer that dynamically modulates the temporal representation at the first stage of cerebellar processing.

## Results

High-power confocal imaging of vermal and floccular slices was used to assess the regional density of PC collateral synapses and UBCs, and to detect putative synaptic connections between PCs and UBCs. PC presynaptic boutons were labelled in Pcp2-Cre x synaptophysin-tdTomato mice. Weaker labelling was evident in PC dendrites and somata (red, Figure 1A). Inhibitory synapses were labelled with VGAT immunofluorescence (green, Figure 1A). PC synapses were automatically identified with a custom deep neural network using VGAT and Synaptophysin-tdTomato immunofluorescence (Figure 1 - figure supplementary 1, see Methods). For visualization, annotated PC synapses were colored in cyan, and the PC and molecular layers are shown in grey based on the synaptophysin-tdTomato signal (Figure 1B). In the same slices, mGluR1 immunofluorescence labelled UBCs and PCs. For display purposes, mGuR1 labelling in the PC and molecular layers was masked to highlight UBCs in the granular layer (magenta, Figure 1. C-K).

**Figure 1.**
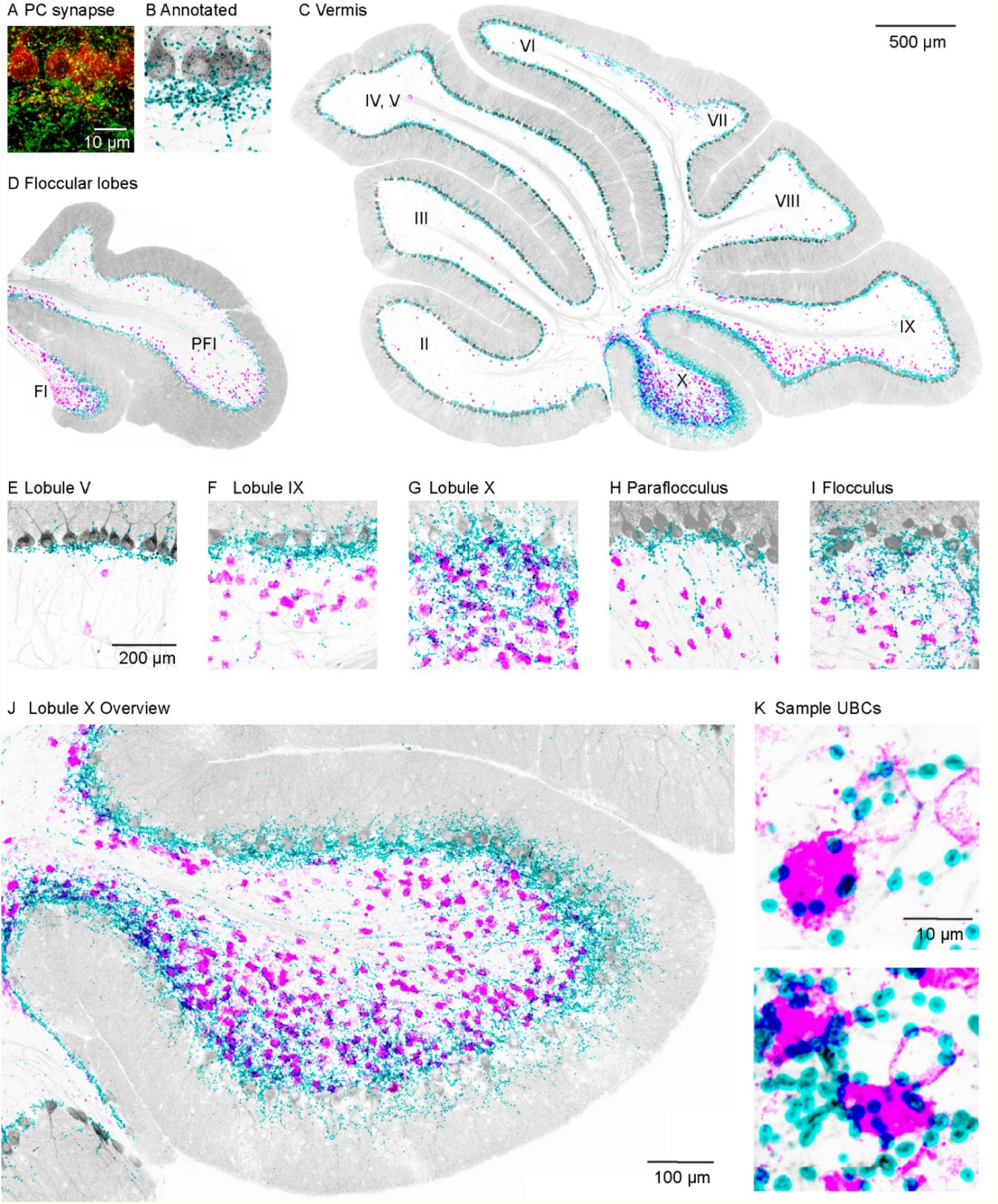
PC collateral synapses colocalized with mGluR1+ UBCs in the vestibular lobules. A. Maximal intensity projection of a 15µm confocal z-stack images of PC collateral synapses co-labeled by synaptophysin-tdTomato (red) and VGAT (green). B. Annotated PC collateral synapses (cyan) and the synaptophysin-tdTomato signal (grey) C. A vermal slice of cerebellum analyzed as in B with synapses labeled in cyan, synaptophysin-tdTomato in grey and the mGluR1 labeled UBCs dendritic brush in magenta. D. Same analysis on a coronal slice of a floccular lobe E-I. Zoomed in view of selective lobules showing variable degrees of colocalization between PC synapses and UBCs. J. Expanded view of Lobule X. K. Two sample UBCs showing clear examples of collateral synapse onto the brush.

The densities of UBCs and granular layer PC synapses exhibited considerable regional variability. The highest densities of both UBCs and granular layer PC synapses were found in vestibular regions (lobules IX, X, FL and PFL see Figure C, D and F-I) and in the oculomotor vermis (lobule VIb and VII, Figure 1C). Much lower densities were observed in anterior cerebellar cortex (lobule I-V, and VIa, see Figure1 C, E) and lobule VIII. In lobule X, there was a particularly high density of PC collateral synapses and UBCs (Figure 1J). High-power images of lobule X suggest that PC synapses directly terminate on the brushes of many UBCs (Figure 1K).

To determine whether PCs directly inhibit UBCs, we optically stimulated PC axons in Pcp2-Cre x ChR2 mice while recoding from UBCs (Figure 2A). We performed these experiments in lobule X of acute slices in the presence of AMPA, NMDA, glycine and GABA_B_ receptor blockers and a high chloride internal solution. UBCs exhibit diverse responses to MF activation, ranging from excitation mediated by AMPARs and mGluR1, to inhibition mediated by mGluR2, with intermediate cells having more diverse responses mediated by a combination of mGluR1 and mGluR2 (Guo et al., 2020). Electrophysiological studies have shown that fast inhibition in UBCs is mediated by a combination of glycine and GABA_A_ receptors (Dugué et al., 2005; Rousseau et al., 2012), but that mGluR1-lacking UBCs lack GABA_A_ receptors (Rousseau et al., 2012). Consistent with these observations, molecular profiling of UBCs using single cell RNAseq (Kozareva et al., 2020) indicate that α1, β2, β3, and γ2 subunits of GABA_A_ receptors have a gradient of expression levels and are selectively enriched in mGluR1 positive UBCs (Figure 2 – figure supplement 1B), but that glycine receptors are broadly expressed by UBCs (Figure 2 – figure supplement 1C). This opens the possibility that the molecular identity of UBCs could determine whether they receive PC feedback. We therefore characterized UBC responses to brief puffs of glutamate (Figure 2B, C left) prior to measuring optically evoked synaptic responses (Figure 2B, C right). Glutamate evoked a continuous range of metabotropic responses across UBCs with variable degree of excitation vs. inhibition, as is apparent in the sorted heatmap of glutamate-evoked response (Figure 2D). Optical stimulation evoked currents in a subpopulation of these cells (Figure 2B right, Figure 2D asterisk marks, 17/41 cells) but not in others (Figure 2C, right, 24/41 cells). GABA_A_R antagonist gabazine eliminated these light-evoked synaptic currents (Figure 2E). We compared the average responses to a glutamate puff for UBCs that had light evoked GABA_A_ currents with those that did not. Cumulative histograms for glutamate-evoked charge transfer (Figure 2F), and the average glutamate-evoked current for UBCs with or without PC feedback (Figure 2G), show that PC GABA_A_ feedback preferentially targets UBCs that are excited by glutamatergic inputs.

**Figure 2.**
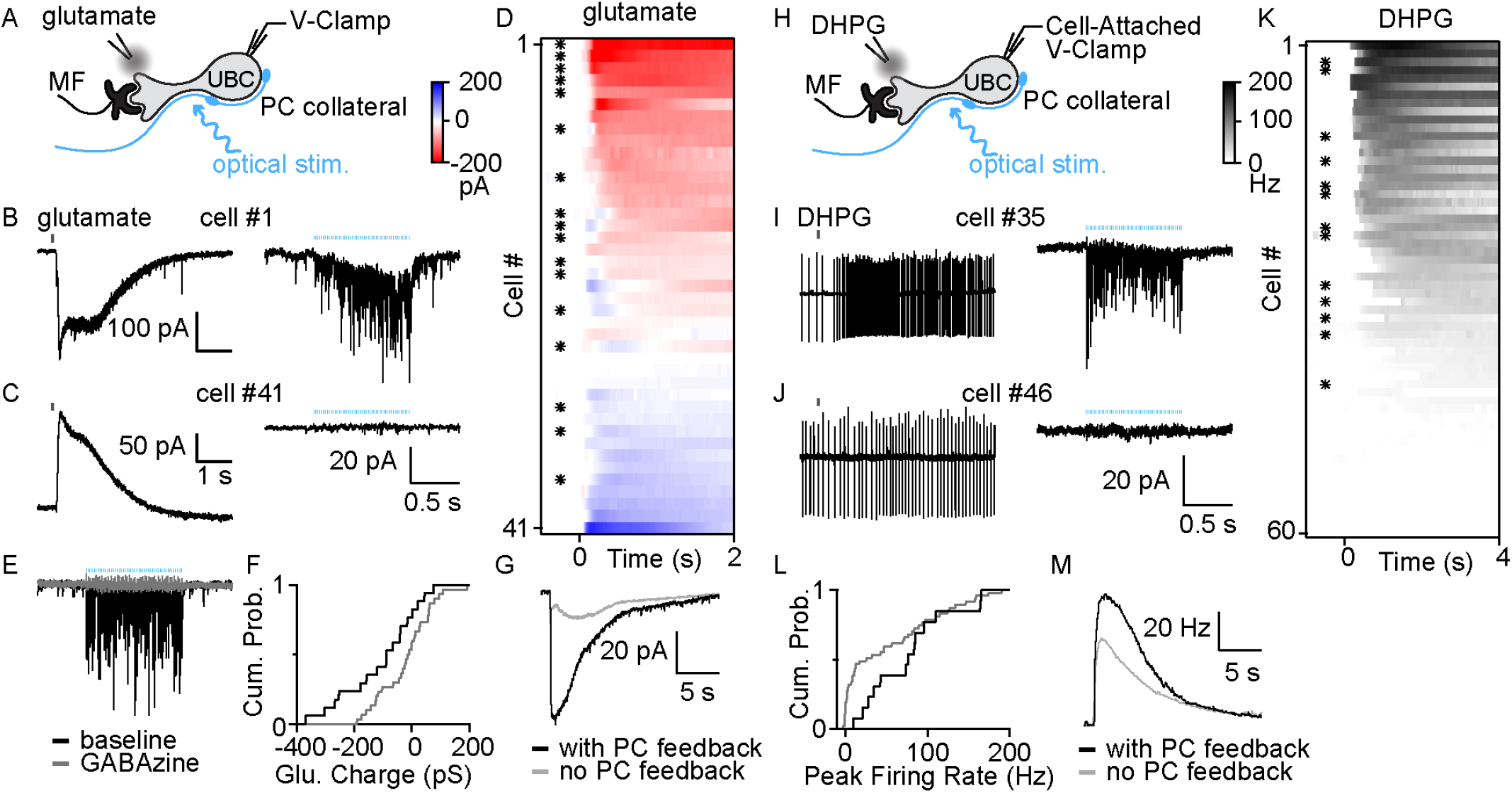
PC collaterals preferentially inhibit mGluR1+ UBCs via fast GABA_A_R-mediate feedback. A. Schematic showing a mossy fiber (MF) and a PC collateral innervating a UBC. A whole-cell electrode was used to voltage clamp the UBC, and responses were measure for either a glutamate puff applied with a nearby electrode, or to optical stimulation of PC collaterals. B. Whole-cell recordings of glutamate evoked currents (left) and to optically evoked PC inhibition (right) are shown for cell #1 of (D). C. As in B, but for cell #41 of (D). D. Summary of cells in which responses to glutamate puffs were measured, with red corresponding to an excitatory inward current and blue corresponding to an inhibitory outward current. Cells with light-evoked inhibitory synaptic current are indicated with a black dot. E. Light-evoked synaptic currents recorded in baseline (black) and after the addition of the GABA_A_ receptor antagonist gabazine (grey). F. Normalized cumulative distributions of charge evoked by a glutamate puff for cells that either had light evoked responses (black), or that did not (grey). G. Average currents evoked by a glutamate puff for cells that had PC feedback (black) and that did not have PC feedback (grey). H. Schematic as in A, but for application of the mGluR1 agonist DHPG. DHPG-evoked increases in firing were measured with a cell-attached electrode, and then optically-evoked synaptic currents were measured with a whole-cell voltage clamp recording. I. DHPG-evoked increases in firing (left) and light-evoked synaptic currents (right) are shown for cell #35 in K. J. Same as I but for cell #45 in K. K. Summary of cells in which firing evoked by DHPG puffs was quantified. Cells with light-evoked inhibitory synaptic current are indicated with a black dot. L. Normalized cumulative distributions of charge evoked by a DHPG puff for cells that either had light evoked responses (black), or that did not (grey). M. Average currents evoked by a DHPG puff for cells that had PC feedback (black) and that did not have PC feedback (grey).

To further characterize the properties of UBCs that receive PC feedback, we categorized UBCs based on their responses to the mGluR1 agonist DHPG prior to characterizing light-evoked synaptic responses (Figure 2H). DHPG puffs evoked spiking in a subset of UBCs (Figure 2I, J left, K 45/60). After recording the DHPG-evoked increased in firing with a cell-attached electrode, we obtained a whole-cell configuration and measured light-evoked responses. 13/45 DHPG responsive UBCs also had significant light-evoked responses (Figure 2I, J, right, K). We then grouped UBCs by the presence (Figure 2I right and asterisk in K) or absence of PC feedback (Figure 2J, right). Cumulative DHPG-evoked peak firing in UBCs with PC feedback is shifted towards greater response than those without feedback (Figure 2L). Furthermore, the average DHPG-evoked instantaneous firing in UBCs with PC feedback is also slightly bigger than those without (Figure 2M). Cumulative histograms for DHPG-evoked firing rate increases in UBCs (Figure 2L), and the average DHPG-evoked firing rate increases for UBCs with or without PC feedback (Figure 2M), show that PC GABA_A_R-mediated feedback preferentially targets UBCs with larger mGluR1-mediated excitation.

We also directly recorded from connected PC-UBC pairs (Figure 3, n = 2). An on-cell recording allowed us to noninvasively monitor PC spiking while recording synaptic responses in a UBC. Many IPSCs in the UBC were timed to PC spikes (Figure 3A top), as is readily appreciated in spike-triggered averages of UBC IPSCs (Figure 3B). The IPSC latency was 1.5±0.4ms (Figure 3C, D, F) and the distribution of response amplitude was approximately Gaussian (Figure 3E). While the response potency is large (Figure 3F), there was a high failure rate of approximately 85% (Figure 3F), which is comparable to PC to GrC synapses (87±2%) (Guo et al., 2016) and higher than PC to PC synapses (56±7%) (Witter et al., 2016). This suggests that phasic GABA_A_R-mediated PC-UBC synapses convey noisy and unreliable output to input layer feedback.

**Figure 3.**
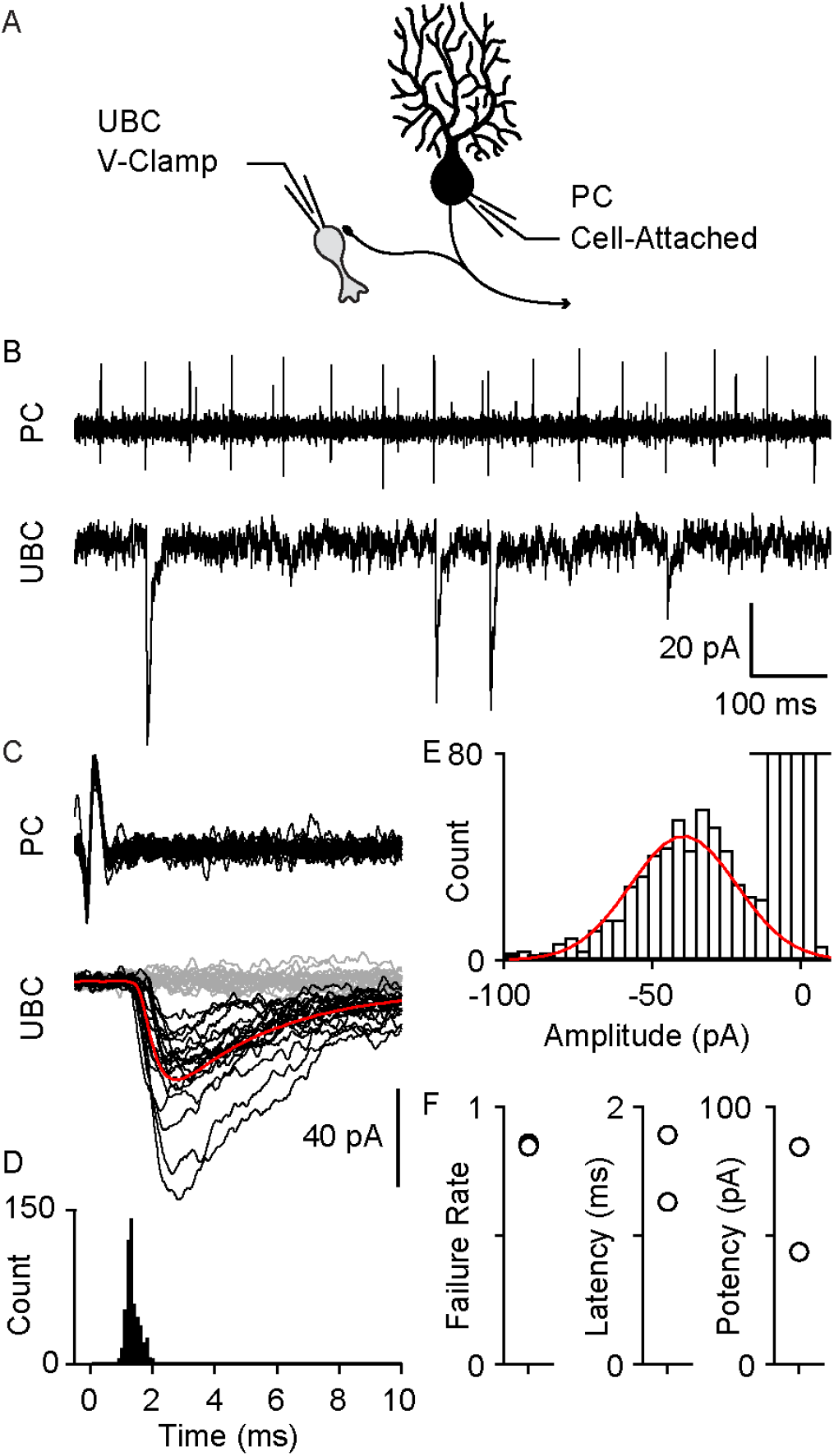
Paired recordings of PC to UBC synaptic connections. A. Schematic of the paired recording configuration B. A representative pair showing a cell-attached PC recording (top) and whole-cell voltage-clamp recording of a synaptically connected UBC (bottom) C. Same two cells showing time-aligned PC spikes (top) and associated IPSCs in UBCs (bottom), with successes in black, failures in grey, and average response from success trials in red. D. Histogram of IPSC latency (bin size = 0.1 ms). E. Histogram of IPSC amplitude (bin size = 4 pA) and Gaussian fit over success trials (red). F. Failure rate (left), latency (middle) and potency (left) PC to UBC synapse (n=2 cells)

One of our goals was to determine if PCs also activate GABA_B_Rs and thereby tonically inhibit UBCs. Previous studies established that only mGluR1-positive UBCs contain GABA_B_ receptors (J.-A. Kim et al., 2012). This is consistent with scRNA_seq_ analysis of UBCs (Figure 2 – figure supplement 1A and D). GABA_B_Rs are comprised of GB1 and GB2 subunits (Gassmann & Bettler, 2012) that are encoded by *Gabbr1* and *Gabbr2*. UBCs exhibit a gradient of expression of *Gabbr2* that is similar to *Grm1* (mGuR1), although *Gabbr1* has a less pronounced gradient of expression (Figure 2 – figure supplementary 1D). We therefore restricted our recordings to cells that responded to DHPG, and that were therefore mGluR1 positive. We found that in whole-cell recordings the currents evoked by GABA_B_R agonists washed out within minutes (Figure 4 – figure supplement 1A and B grey). This washout was accompanied by an increase in the leak current (Figure 4 – figure supplement 1C). In contrast, in perforated patch recordings GABA_B_ responses were large and stable, and leak currents were also stable (black, Figure 4 – figure supplement 1). These experiments established that perforated patch recordings facilitate the measurement of PC-evoked GABA_B_ responses.

We examined the PC-evoked GABA_B_ responses in UBCs with an approach similar to that used to record GABA_A_ responses, except that we blocked GABA_A_ receptors, used perforated patch recordings, and elevated PC firing for 2s to activate the GABA_B_ receptors on a longer time scale (Figure 4A). We stimulated the entire lobule X with graded light levels to approximately double and triple PC firing rates (Figure 4B-D). Optical stimulation evoked outward currents in all UBCs (Figure 4E-G, 13/13). GABA_B_R-mediated inhibition was slow (Figure 4F, rise time constant = 583±6ms, decay time constant = 533±5ms). The GABA_B_R antagonist CGP eliminated these light-evoked responses (Figure 4K). The average inhibition of 3.8±2.2pA was sufficiently large to strongly suppress or entirely shut down spontaneous UBC firing (Figure 4H-J in 9/10 spontaneous firing UBCs). These findings suggest that PC firing rates dynamically regulates the activation of GABA_B_Rs and by extension the spontaneous firing of most mGluR1 positive UBCs. With this GABA_B_R-dependent mechanism, PCs inhibit a much larger fraction of mGluR1 positive UBCs (90∼100% Figure 4G, 13/13 DHPG responding cells and Figure 4J, 9/10 spontaneous firing), compared to GABA_A_R-dependent inhibition (29%, Figure 2K, 13/45 DHPG responding cells).

**Figure 4.**
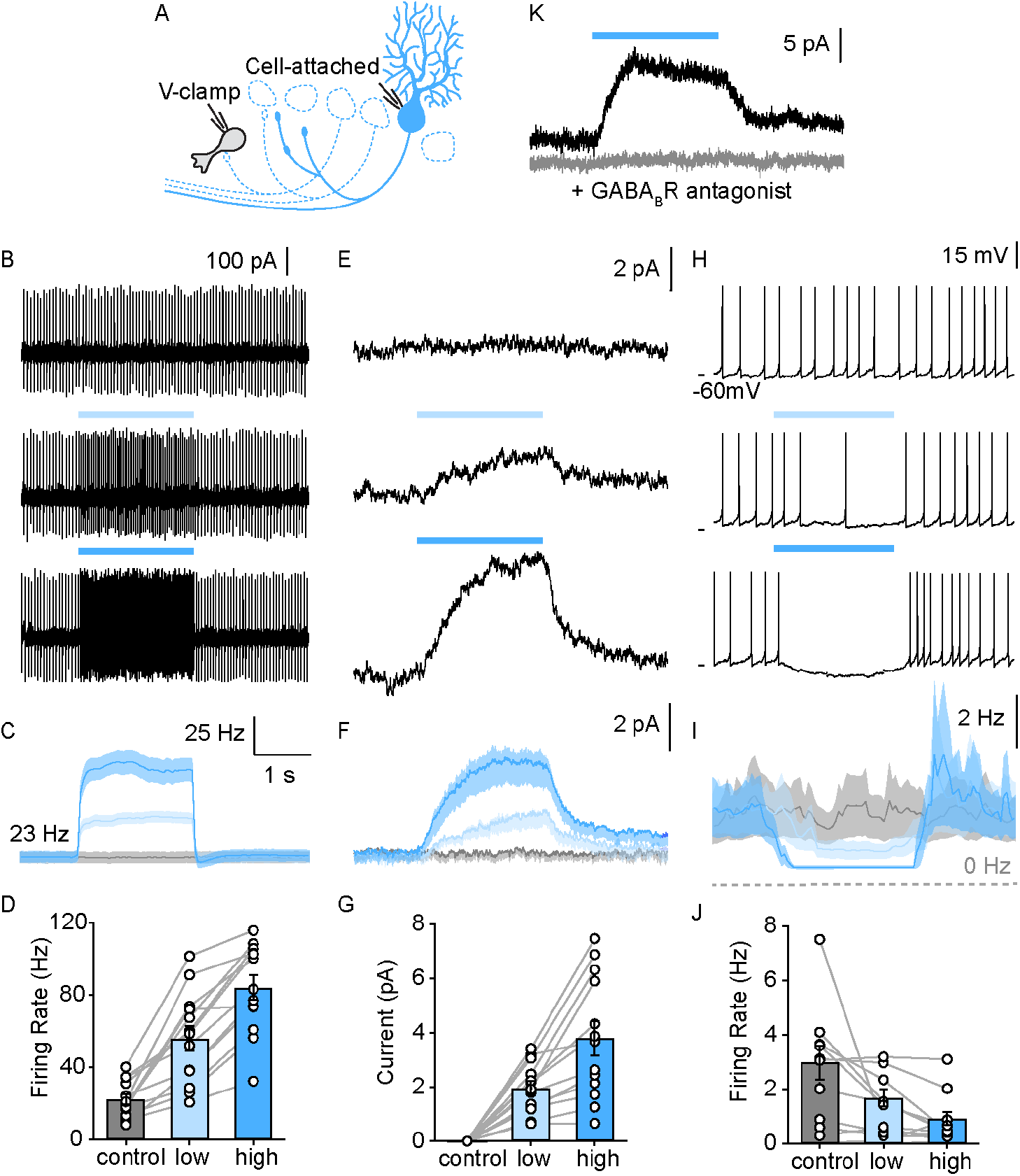
PC collaterals provide slow GABA_B_ receptor mediated inhibition onto lobule X UBCs. A. Schematic of full-field optical stimulation for modulating PC firing rate in lobule X B. Cell-attached recording of spontaneous PC spikes for no (top), low intensity (middle) and high intensity (bottom) full-field optical stimulations. C. Average instantaneous firing rate of PCs with no (grey), low (light blue) and high (blue) intensity stimulations (n = 13) D. Summary of PC firing rate with no (grey), low (light blue) and high (blue) intensity light-stimulations of PCs (n = 13) E. Voltage-clamp recordings in UBC revealed a slow inhibitory response that is modulated by PC firing rate, same optical stimulation conditions as in B F. Average evoked current of UBCs during stimulations as in E (n= 13) G. Summary of inhibitory current amplitude in UBCs with no (grey), low (light blue) and high (blue) intensity light-stimulations of PCs (n = 13) H. Full field activation of PCs suppressed the frequency of UBC firing. I. Average instantaneous firing rate of UBCs with no (grey), low (light blue) and high (blue) intensity stimulations (n = 13) J. Summary of firing rate changes in spontaneous firing UBCs with no (grey), low (light blue) and high (blue) intensity light-stimulations of PCs (n = 10) K. PC evoked slow outward currents in UBC in the presence of GABA_A_R blockers under voltage-clamp before (black) and after blocking GABA_B_R (grey).

In addition to suppressing spontaneous activity, PC firing and GABA_B_R activation could control the excitability of UBCs and their response to depolarization. To assess the effect of small tonic hyperpolarizing currents on UBC excitability, we examined how such currents influence UBC responses to current steps using perforated patch recordings. In spontaneously firing UBCs, small depolarizing current steps evoked firing that declined to a somewhat reduced steady-state response that was proportional to the current injected (Figure 5A, B and C grey left). A 5pA tonic hyperpolarizing current, drastically altered these responses, with current steps evoking a much larger transient responses and reduced steady-state firing (Figure 5A, B and C black right). A summary of average spiking responses to different current step amplitudes of all UBCs without and with tonic hyperpolarization is shown in Figure 5B and C. In all cells, hyperpolarization increased the difference between peak and steady-state firing rates (Figure 5C, n = 4). We also directly examined the effects of PC firing on UBC excitability. In these experiments, we blocked GABA_A_Rs to isolate the effect of GABA_B_Rs on spiking. Full-field optical excitation of PCs activated GABA_B_R-mediated inhibition and produced qualitatively similar effects on spiking as hyperpolarizing current injections (Figure 5D, blue right). Optical activation of PC feedback through GABA_B_R hyperpolarized UBCs (Figure 5D, blue right), and current steps evoked larger initial responses and smaller steady-state responses (Figure 5E, F blue vs. grey). These counterintuitive effects on spiking, in which a hyperpolarization increases initial responses, are consistent with previously described properties of UBCs (see Discussion). Thus, tonic PC feedback via GABA_B_R-mediated inhibition of UBCs results in temporal sharpening of spiking response to excitatory MF input.

**Figure 5.**
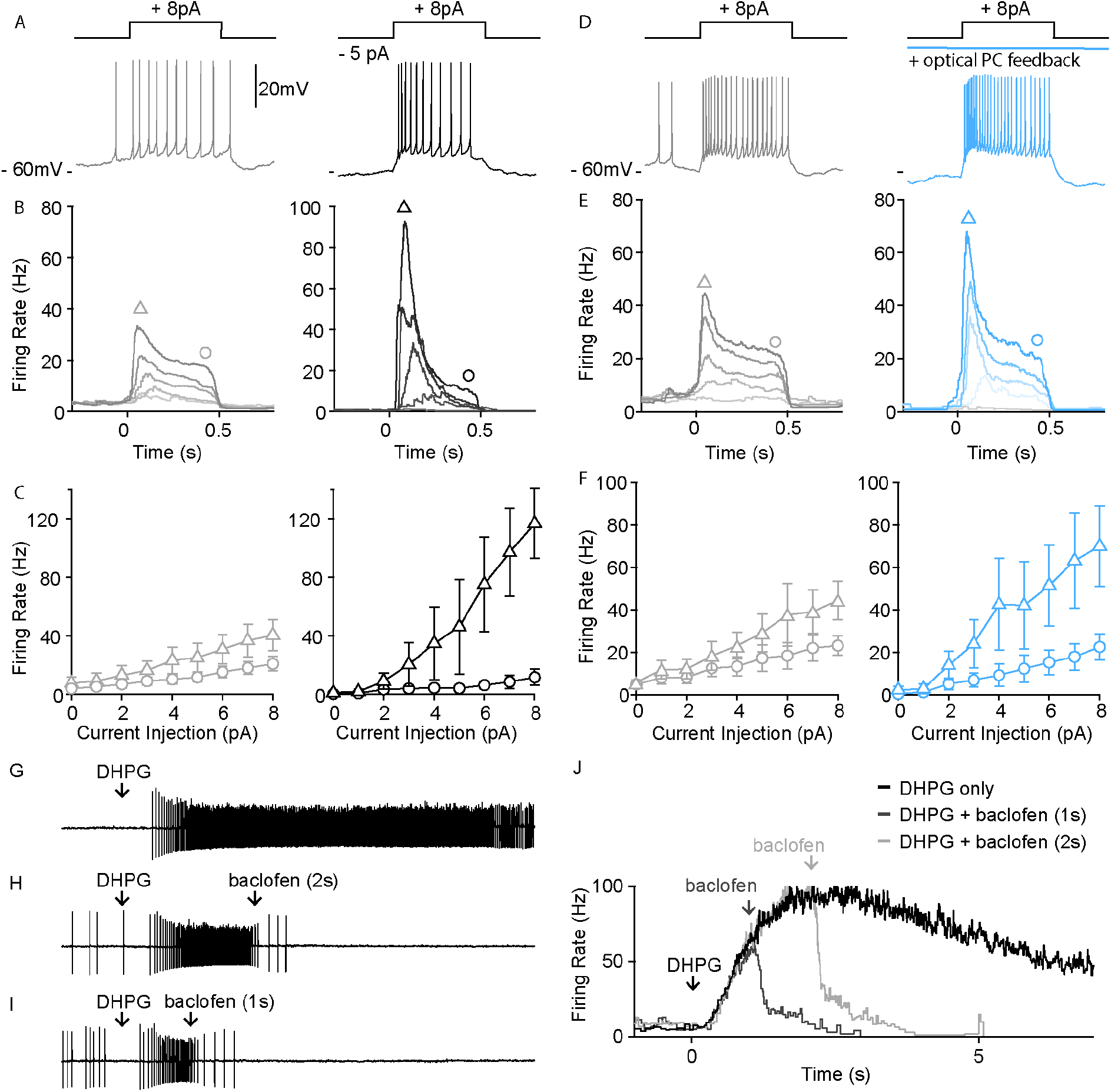
PC-UBC feedback via GABA_B_R sharpens temporal response to current injections. A. Sample perforated current-clamp recording of UBC spiking response to an 8pA current step (500 ms), without (grey, left) and with a hyperpolarizing tonic current (black, right) B. Average instantaneous firing rate to current steps of (0pA, 2pA, 4pA, 6pA, and 8pA, from light to dark shade) without (grey, left) and with a hyperpolarizing tonic current (black, right) C. Summary of peak firing rate (triangle marker, mean ± sem) and steady state firing rate (circle marker, mean ± sem) for the two conditions as in B. D. Same plot as in A but for control (grey, left) vs. optical PC feedback (blue right) E. Same summary plot as in B for control (grey, left) vs. optical PC feedback (blue, right) F. Same summary plot as in C for control (grey, left) vs. optical PC feedback (blue right) G. In a sample UBC pressure-application of an mGluR1 agonist (DHPG 100µM, 10ms) evoked persistent spiking for a few seconds. H. Pressure-applied GABA_B_R agonist (baclofen 250μM, 100ms) 2s following mGluR1 agonist application readily reduced the firing rate of persistent spiking response. I. Same as in B but only 1s following mGluR1 agonist application. J. Average instantaneous firing rate of the sample UBC under mGluR1 agonist application only (black), mGluR1 + GABA_B_R agonist applied 1s apart (dark grey) or 2s apart (light grey).

MF activity elevates firing in the input layer of the cerebellar cortex that in turns alters PC spiking. The PC to UBC feedback described here has the potential to allow PC output to dynamically regulate the firing of mGluR1-expressing UBCs. This idea is illustrated by brief DHPG to crudely mimic MF-evoked activation of mGluR1-expressing UBCs, and baclofen to mimic GABA_B_ receptor activation arising from elevated PC firing (Figure 5G, J). Brief application of baclofen following DHPG resulted in a reduction in evoked spikes (Figure 5H-J). Thus, it is likely that MF input-driven increases of PC firing will feedback to UBCs in the input layer to either suppress the firing or sharpen the temporal kinetics of most mGluR1-expressing neurons. This feedback though the GABA_B_R will be slow and will outlive transient changes in PC firing. In this way, UBC responses *in vivo* are not simply feed forward, but are regulated both tonically and dynamically by PC feedback.

## Conclusions

The PC to UBC synapses described here link two previously described regional specializations of the cerebellar cortex: the high densities of UBCs, and the prevalence of PC synapses within the granular layer. PC to UBC synapses provides a new way for PC outputs to dynamically regulate the temporal transformations by UBC populations within the input layer.

### Target specificity of GABA_A_ and GABA_B_ PC to UBC feedback

PCs inhibit a subset of UBCs. Optical stimulation of PC terminals evoked GABA_A_R-mediated responses in 29% (30/101) of UBCs. Light-evoked PC GABA_A_R-mediated inhibition was much more common in cells in which glutamate evoked a large excitatory current but was also observed in three cases where glutamate evoked inhibitory currents (Figure 2D). This may reflect the graded nature of metabotropic glutamate receptor signaling in UBCs in which outward mGluR1-mediated currents and inward mGluR2 mediated currents can be present in the same cell, and a large mGluR2 component could obscure a small mGluR1 component in some cells (Guo et al., 2020; Kozareva et al., 2020). When we used DHPG to identify mGluR1+ cells, we found that PCs did not inhibit any mGluR1-UBCs (0/15), (Figure 2K). This is in keeping with the observations that mGluR1-UBCs do not respond to either GABA_A_R or GABA_B_R agonists (J.-A. Kim et al., 2012; Rousseau et al., 2012), that they do not express GABA_A_ or GABA_B_ receptor subunits (Rousseau et al., 2012) (Figure 2 – figure supplementary 1), and that PCs do not release glycine (Tanaka & Ezure, 2004). In contrast to PCs that only inhibit a subset of UBCs, glycinergic Golgi cells can inhibit all UBCs, because all UBCs express ionotropic glycine receptors and glycinergic IPSCs are observed in all UBCs (Dugué et al., 2005; Rousseau et al., 2012).

Our findings provide new insights into the previous immunohistochemical characterization of the inhibitory synapses onto UBCs that suggested approximately 50% corelease GABA and glycine and the rest only release GABA (Dugué et al., 2005; Rousseau et al., 2012). Previously it was thought that these exclusively GABAergic synapses were made by non-glycinergic Golgi cells. Our findings suggest that PCs likely constitute most of these synapses.

PCs inhibit UBCs by activating both GABA_A_ receptors and GABA_B_ receptors, and light-evoked GABA_B_ currents are observed in a larger percentage of mGluR1+ UBCs (>90%) than are light-evoked GABA_A_ currents 29% (13/45). This difference in prevalence likely reflects differential sensitivities of GABA_A_ and GABA_B_ receptors. Activation of GABA_A_Rs on UBCs requires high GABA levels (Möhler, 2006), and thus require a direct PC synapse for effective activation. GABA_B_Rs are activated by much lower levels of GABA (Galvez et al., 2000; Kaupmann et al., 1998). Consequently, a direct PC input is not required and GABA released from PC synapses onto neighboring cells can pool and spillover to activate GABA_B_Rs. This also suggests that the combined activity of many PCs could efficiently activate GABA_B_Rs on UBCs and control spontaneous activity.

### Comparisons to properties of PC-GrC feedback

Within the input layer, PCs also directly inhibit GrCs, and there are similarities and differences in the inhibition of these two targets. The failure rate and potency of the fast component of inhibition is similar for PC synapses onto UBCs and GrCs (Guo et al., 2016). Therefore, we hypothesize that the overall effect of fast and stochastic GABA_A_R PC inhibition of UBCs and GrCs is to lower the gain of the input layer. The slow GABA_B_ feedback to UBCs occurs in concert with a slow inhibition of GrCs that is mediated by a very different mechanism (Guo et al., 2016). PCs slowly inhibit GrCs primarily by activating special high-affinity α6 and δ subunit containing GABA_A_ receptors that are specialized to respond to low levels of extracellular GABA (Brickley et al., 1996; Hamann et al., 2002; Kaneda et al., 1995; Wall & Usowicz, 1997). It appears that α6/δ containing GABA_A_Rs in GrCs and GABA_B_Rs in UBCs can both respond to the low GABA levels that reflect the population averaged PC firing rates, and in both cases the kinetics of the responses are slow (312ms in GrCs vs. 580ms in UBCs). The major differences in slow inhibition of UBCs and GrCs are the state change of UBCs and the target dependence of the UBC inhibition, neither of which have been described for PC inhibition of GrCs (Guo et al., 2016)

### Functional Consequences of Fast and Slow PC to UBC inhibition

PC inhibition of UBCs by activation of GABA_A_Rs and GABA_B_Rs likely has distinctive functional consequences. The ionotropic GABA_A_ component is much faster than the metabotropic GABA_B_ component, as is typically the case (Dutar & Nicoll, 1988; U. Kim et al., 1997). The high failure rates of PC-UBC GABA_A_R synapses, in which just one in ten PC spikes results in a post-synaptic response in both UBCs and GrCs, suggest that both first order timing information in the inter-spike intervals in PC output and second order timing information from PC spike synchrony will be lost (Han et al., 2018). Instead, GABA_A_R-mediated feedback can be thought of as a source of noise that is modulated by the PC firing rate. Such noisy synaptic conductance can linearize f-I curves near threshold and decrease response gain (Chance et al., 2002; Mitchell & Silver, 2003).

The effect of the slow GABA_B_ component on UBCs is very different. Slow PC-UBC inhibition represents the average synaptic contribution from multiple PCs, and when the firing rates of these PCs are elevated UBCs are hyperpolarized. Increases in PC firing lead to slow increases in inhibition, and following decreases in PC firing rate this inhibition gradually decreases. Importantly, this alters the state of the UBC. As has been shown previously, hyperpolarization can relieve inactivation of T-type calcium channels (Perez-Reyes, 2003) and lead to UBCs that are more prone to bursting leading to a steeper initial input/output relationship for depolarizing inputs (Figure 5, EF). In this way PC firing rate changes can alter the UBC-mediated transformations in the input layer of the cerebellum.

Slow inhibition can also allow PCs to dynamically regulate the excitability of mGluR1+ cells. MF excitation or inhibition of UBCs will modulate the firing of PCs and these firing rate changes will feed back to mGluR1+ UBCs to alter their firing. Similar to the slow GABA_A_-mediated PC-GrC feedback that is also present in vestibular regions, the slow GABA_B_ component may also confer the network certain advantages in terms of stable representation and learning over longer time scales.

## Materials and Methods

### Immunohistochemistry and confocal imaging

Pcp2-Cre (Jackson Laboratory, 010536) x synaptophysin-tdTomato (Ai34D, Jackson Laboratory, 012570) mice were anesthetized with intraperitoneal injections of ketamine/xylazine/acepromazine mixture at 100,10 and 3 mg/kg and transcardially perfused with 4% PFA in phosphate-buffered saline solution. The brain was removed from the skull and post-fixed overnight in 4% PFA. The cerebellar vermis and flocculus were dissected out and embedded in 6% low melting agarose before slicing. Sagittal vermal slices and coronal floccular slices at 50μm were obtained from two animals using Leica VT1000S vibratome (Leica Biosystems, Buffalo Groves, IL). For immunolabeling of inhibitory synapses and the unipolar brush cells, slices were permeabilized (0.4% Triton X-100 and PBS) for 30 mins and blocked for 1 hour at room temperature (0.4% Triton X-100, 4% normal goat serum) and incubated overnight at 4°C with primary antibodies (Guinea-pig anti-VGAT, Synaptic Systems 131004, 1μg/mL, 1:200 and mouse anti-mGluR1, BD Pharmingen,0.5mg/mL, 1:800 in 0.2% Triton X-100 and 2% normal goat serum). The following day the slices were washed three times for 10 minutes in PBS and incubated with secondary antibodies overnight at 4°C (Goat-anti-Guinea-pig-Alexa Fluor 488, Abcam ab150185, 1:500 and Goat-anti-mouse Alexa Fluor 647, Invitrogen A32728, 1:500). Slices were mounted with a #1.5 coverslip using ProLong™ Diamond Antifade Mountant without DAPI (Thermo Fisher Scientific, Waltham, MA).

Z-stack images of entire vermal and floccular slices were obtained on the Olympus FV3000 confocal microscope using the Multi-Area Time-Lapse Software module. A 60X oil immersion objective with 1.42 NA was used and the x,y,z resolutions were 0.212 μm, 0.212 μm, and 0.906 μm respectively. Each z-step was 1μm and the entire stack was 20μm in thickness.

### Image analysis

The PC synapse was detected using a modified deep neural network that is based on the U-Net (Ronnerberger, 2015). The network was fed in all three imaging channels including the mGluR1 labeling. The rationale being that while colocalization of Synaptophysin and VGAT was sufficient for synapse detection, mGluR1 provided additional contextual information for annotation in the molecular layer as it densely labeled PC dendrites. The network was trained using soft-dice loss on 100 annotated images and validated with an 80/20 train-test-split. We found that ensembling with 10 independently trained networks qualitatively improved the annotation. The true positive rate of synapse detection was at 95.5% and the false-positive rate was 10.11%. We note however that the false positive rate was likely inflated as the human annotator sometimes missed difficult to spot synapses (Supplementary Figure 1. Yellow arrows). For illustrating the UBCs, the mGluR1 signals in the molecular layer were manually cropped in the final composite image.

### Slice preparation for electrophysiology

Adult (P30-P40) C57BL/6 or Pcp2-Cre x ChR2-EYFP (Ai32, Jackson Laboratory,024109) mice were first anesthetized with an intraperitoneal injection of ketamine/xylazine/acepromazine mixture at 100,10 and 3 mg/kg and transcardially perfused with an ice-cold choline slicing solution consisting (in mM): 110 choline Cl, 7 MgSO_4_, 2.5 KCl, 1.2 NaH_2_PO_4_, 0.5 CaCl2, 11.6 Na-ascorbate, 2.4 Na-pyruvate, 25 NaHCO_3_ and 25 glucose equilibrated with 95% O_2_ and 5% CO_2_. The cerebellum was dissected, mounted against an agar block, and submerged in the choline solution during slicing. Sagittal slices of the vermis were obtained using a Leica VT1200S vibratome and allowed to recover for 30 minutes at 33°C in artificial cerebral spinal fluid (ACSF) consisting of (in mM): 125 NaCl, 26 NaHCO_3_, 1.25 NaH_2_PO_4_, 2.5 KCl, 1 MgCl_2_, 1.5 CaCl_2_, and 25 glucose (pH 7.4, osmolarity 315) equilibrated with 95% O_2_ and 5% CO_2_. The incubation chamber was then removed from the warm water bath and kept at room temperature for recording for up to 6 hours.

### Electrophysiology

Recordings were performed at 34-36°C in ASCF containing 5μM NBQX, 2μM R-CPP and 1μM strychnine set to a flow rate of 2mL/min. Visually guided recording of UBCs were obtained under a 60X objective with differential interference contrast (DIC) imaging on an Olympus BX51WI microscope. Identity of the UBCs was also verified with fluorescent dye after recording (Alexa Fluor 594, 100 μM). Borosilicate patch pipette (3-5MΩ) containing either a KCl internal (in mM: 140 KCl, 4 NaCl, 0.5 CaCl_2_, 10 HEPES, 4 MgATP, 0.3 NaGTP, 5 EGTA, and 2 QX-314, pH adjusted to 7.2 with KOH) or a K-methanesulfonate internal (in mM: 122 K-methanesulfonate, 9 NaCl, 9 HEPES, 0.036 CaCl_2_, 1.62 MgCl_2_, 4 MgATP, 0.3 Tris-GTP, 14 Tris-creatine phosphate and 0.18 EGTA, pH7.4) was used for whole-cell recordings of GABA_A_ or GABA_B_ receptor-mediated current, respectively. A junction potential of -8mV was corrected for the K-Methansulfonate internal during recording. For cell attached UBC recordings, ACSF was used as the internal solution. For perforated-patch recording, an internal containing (in mM) was used: 100 K-methanesulfonate, 13 NaCl, 2MgCl_2_, 10 EGTA, 1CaCl_2_, 10 HEPES, 0.1 Lucifer Yellow and 0.25 amphotericin B, pH 7.2 with KOH. A junction potential of +7mV was corrected online. For recordings of the GABA_B_R-mediated current, 10μM SR95531/gabazine was included in the bath to isolate the metabotropic inhibition. For pair PC-UBC recording, a whole-cell UBC recording with KCl was obtained first. Then 10∼20 PCs in lobule X were screened with cell-attached recordings using a large pipette (1∼2MΩ) containing ACSF.

### Pharmacology

For identification of UBC subtypes in either whole-cell or perforated recordings, ACSF containing glutamate (1mM, 50ms) or DHPG (100 µM, 20ms) was pressure-applied at 5psi through a borosilicate pipette with a Picospritzer™ III (Parker Hannifin, Hollis, NH). For the sequential mGluR1 and GABA_B_R activation experiment (Figure 5G-J), concentrations and durations of pressure-applications are 100 µM, 50ms for DHPG and 250 µM, 100ms for baclofen. The same baclofen concentration and duration were used for testing the stability of GABA_B_R-mediated current (Figure 4 – figure supplementary 1). For wash-in experiments, solution exchange was controlled via ValveLink8.2 Controller (AutoMate Scientific, Berkeley, CA). Gabazine (SR95531, 5μM) was used for blocking GABA_A_R and CGP (1μM) was used for blocking GABA_B_R.

### Optogenetics

Slices from Pcp2-Cre x ChR2-EYFP were kept in the dark before recording. A laser source (MBL-III-473-50mW, Optoengine) was fiber coupled to the excitation path of the microscope. For over bouton stimulation, brief (0.5ms) high intensity (160mW/mm^2^) light pulses were delivered under the 60X objective and focused down to a 50μm spot. For the optical modulation of PC spontaneous firing, much lower intensities (10 μW/mm^2^ or 25 μW/mm^2^) of light were delivered through the 10X objective over the entire lobule X (∼1mm spot) for 2 seconds. To ensure the quality of the slice, we recorded many PCs before the experiment to check for spontaneous activity and to ensure that these low-intensity stimulations did not result in PCs bursting. Only healthy slices were used for subsequent UBC recordings.

### Data Acquisition and Analysis

Electrophysiology experiments were performed with a MultiClamp 700B amplifier (Molecular Device) and controlled by mafPC (Matthew Xu-Friedman, SUNY Buffalo) in Igor Pro 7 (WaveMetrics). Data were filtered at 4kHz in MultiClamp and digitized at 50kHz by InstruTECH ITC-18 (Heka Instrument Inc.). Analysis of DHPG evoked spiking in UBC was done via peak detection in MATLAB. sIPSC timing was determined by time of 5% to peak threshold crossing. The timing of the PC spike for determining the latency of synaptic transmission was determined by the first derivative of the action potential. The time constant for the slow GABA_B_R-mediated current was obtained using a single exponential fit. Data are reported as mean ± sem.

## Acknowledgments

This work was supported by a NIH grant R35NS097284 to W.G.R. Imaging was performed in the Vision Core and NINDS P30 Core Center (NS072030) to the Neurobiology Imaging Center at Harvard Medical School with the help of M. El-Rifai (performed RNA scope®) and M. Ocana. We thank Laurens Witter for his insights in the early stages of this project, and Vincent Huson for feedback on the manuscript.

## Author Contributions

C.G., and W. R. conceived the experiments. C.G. and S.R. both did experiments for Figure 2 H-M. C.G. performed all other experiments. C.G. and M.N. worked together on the image analysis and preparation of Figure 1 and Figure 1 – figure supplement 1. C.G. conducted all other analyses and made illustrations. C.G. and W.G.R. wrote the original draft of the manuscript and C.G., SR and W.G.R revised the manuscript. We thank Laurens Witter for his insights in the early stage of this project, and Vincent Huson for his comments on the manuscript.

---

**Figure 1 – figure supplement 1.**
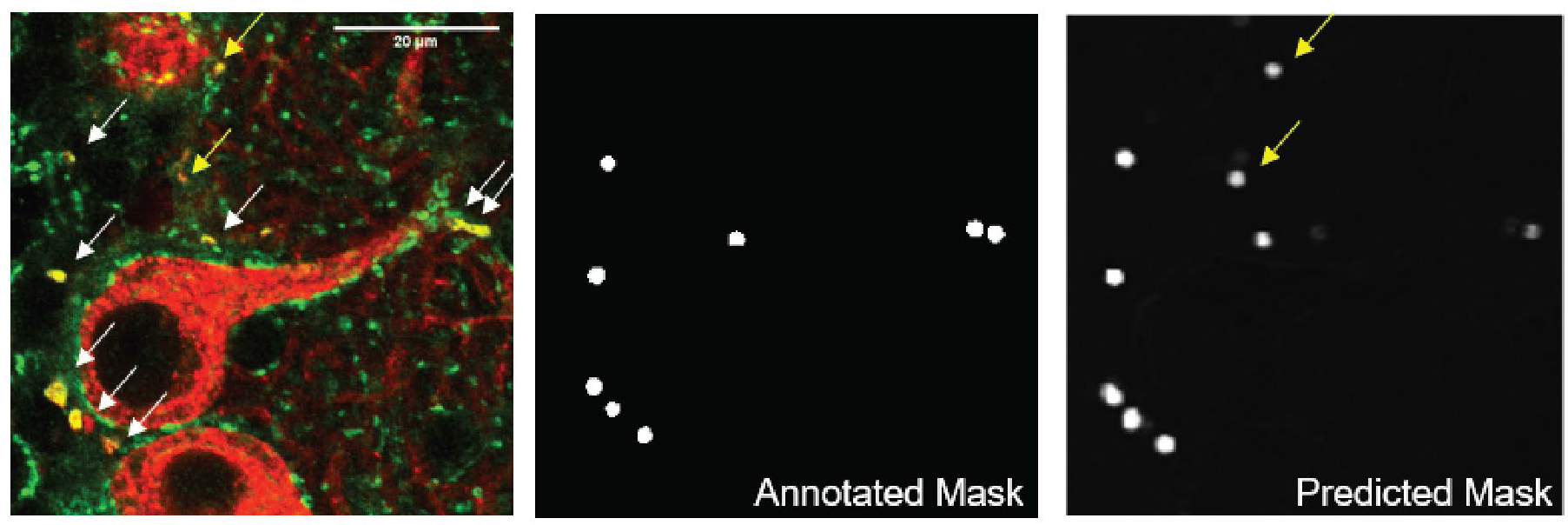
Automatic synapse detection using a convolutional neural network. A. Raw image of PC collateral synapses co-labeled by synaptophysin-tdTomato (red) and VGAT (green). mGluR_1_labeling (not shown) was also used as input to the network as it provided additional contextual information about the location of synapses, which aided automated annotation. Arrowed marked the synapses that annotator picked up (white) and missed (yellow), scale bar = 20 µm. B. Annotated mask used for training the neural network C. Network output after training. The network picked up several synapses that the annotator missed (yellow arrows)

**Figure 2 – figure supplementary 1.**
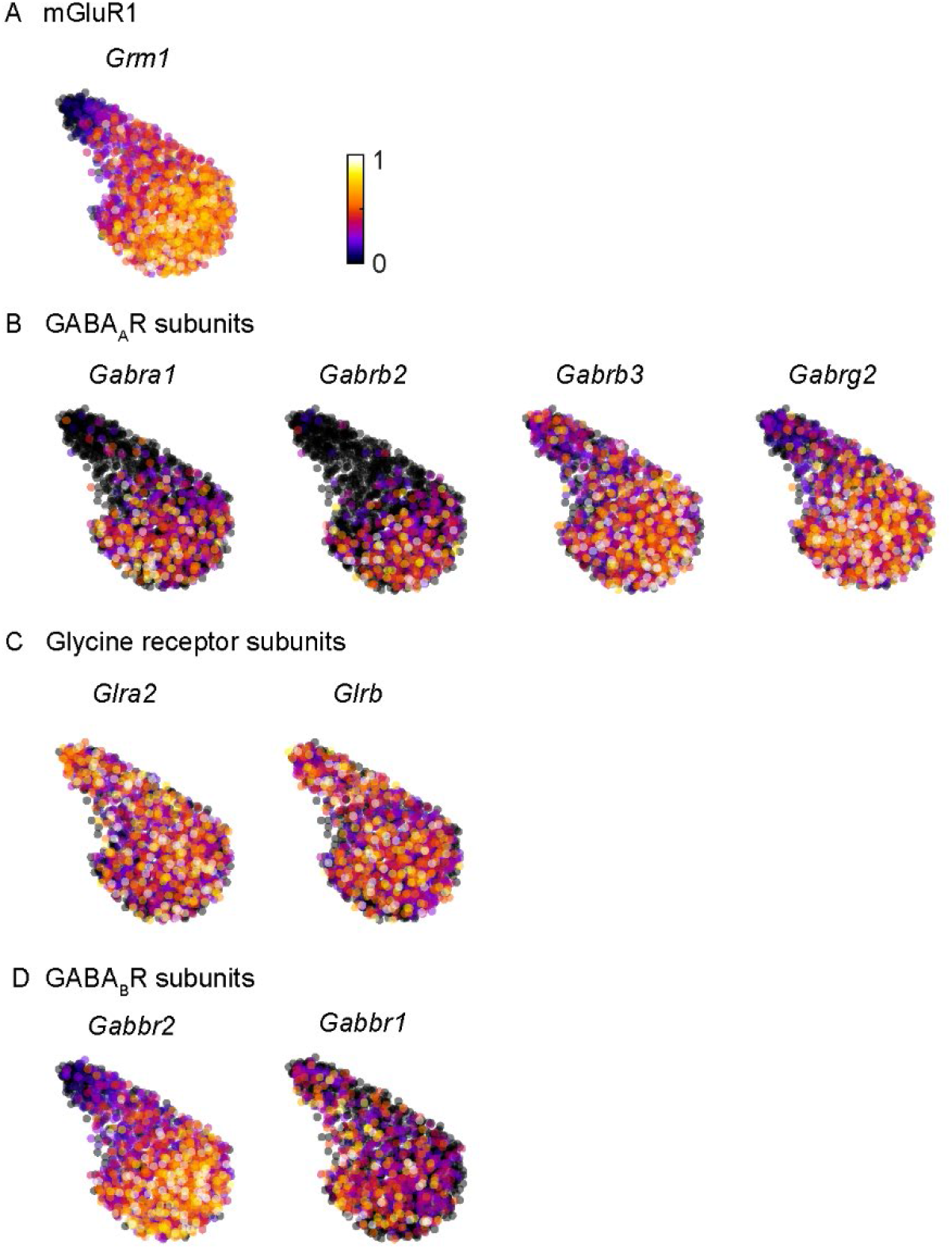
GABA_A_ and GABA_B_ receptor subunits are preferentially expressed in mGluR1 positive UBCs. A. Normalized expression of mGluR1 (*Grm1*) in UBCs with UMAP embedding B. Same plot for GABA_A_ receptor subunits α1, β2/3 and γ2 (*Gabra1, Gabrb2*/*3* and *Gabrg2*) with significant expression in UBCs. C. Same plot for glycine receptor subunits α2 and β (*Glra2* and *Glrb*) D. Same plot for GABA_B_ receptor subunits 1 and 2 (*Gabbr1* and *Gabbr2*)

**Figure 4 – figure supplement 1.**
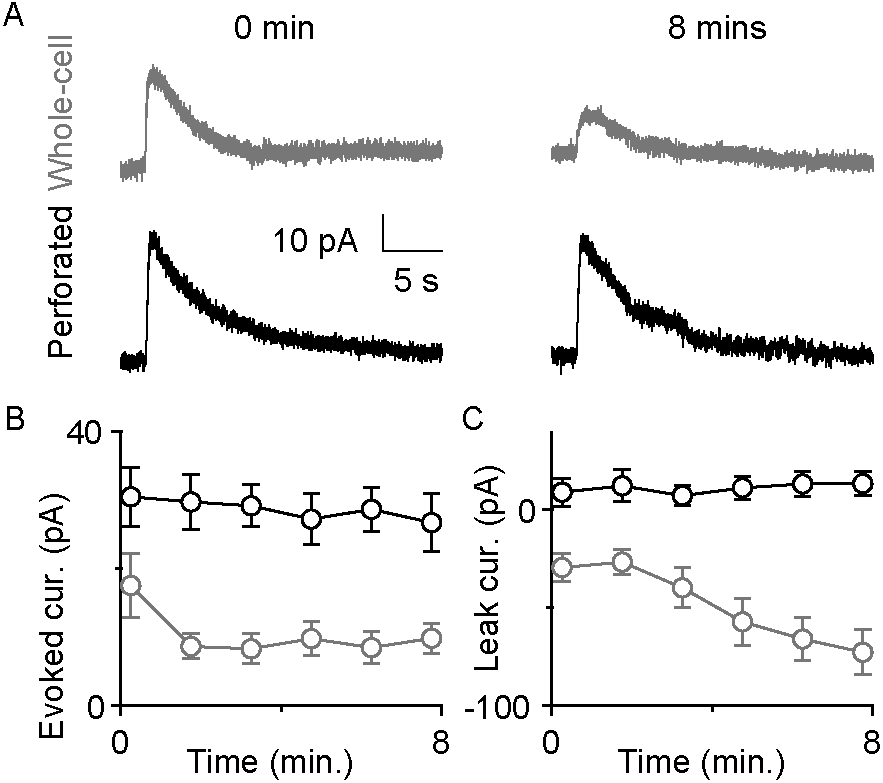
Perforated patch recordings provide stable responses to a GABA_B_-receptor agonist. A. Sample whole-cell (top, grey) and perforated (bottom, black) responses to baclofen puffs immediately after break-in (left) and 8 minutes later (right column) B. Baclofen response amplitudes measured with perforated patch (black) and whole-cell (grey) recordings. C. Magnitude of leak currents measured with perforated patch (black) and whole-cell (grey) recordings.

